# Comprehensive analysis of chromothripsis in 2,658 human cancers using whole-genome sequencing

**DOI:** 10.1101/333617

**Authors:** Isidro Cortés-Ciriano, June-Koo Lee, Ruibin Xi, Dhawal Jain, Youngsook L. Jung, Lixing Yang, Dmitry Gordenin, Leszek J. Klimczak, Cheng-Zhong Zhang, David S. Pellman, Peter J. Park, on behalf of the PCAWG Structural Variation Working Group, the ICGC/TCGA Pan-Cancer Analysis of Whole Genomes Network

**Affiliations:** Department of Biomedical Informatics, Harvard Medical School, Boston, Massachusetts, USA; Ludwig Center at Harvard, Boston, MA 02115, USA; Centre for Molecular Science Informatics, Department of Chemistry, University of Cambridge, Lensfield Road, Cambridge CB2 1EW, United Kingdom; School of Mathematical Sciences and Center for Statistical Science, Peking University, Beijing 100871, China; The Ben May Department for Cancer Research and Department of Human Genetics, The University of Chicago, Chicago, IL 60637; Genome Integrity and Structural Biology Laboratory; Integrative Bioinformatics Group, National Institute of Environmental Health Sciences, US National Institutes of Health, Research Triangle Park, North Carolina, USA; Department of Pediatric Oncology, Dana-Farber Cancer Institute, Boston, Massachusetts 02215, USA; Howard Hughes Medical Institute and Department of Cell Biology, Harvard Medical School, Boston, Massachusetts 02115, USA

## Abstract

Chromothripsis is a newly discovered mutational phenomenon involving massive, clustered genomic rearrangements that occurs in cancer and other diseases. Recent studies in cancer suggest that chromothripsis may be far more common than initially inferred from low resolution DNA copy number data. Here, we analyze the patterns of chromothripsis across 2,658 tumors spanning 39 cancer types using whole-genome sequencing data. We find that chromothripsis events are pervasive across cancers, with a frequency of >50% in several cancer types. Whereas canonical chromothripsis profiles display oscillations between two copy number states, a considerable fraction of the events involves multiple chromosomes as well as additional structural alterations. In addition to non-homologous end-joining, we detect signatures of replicative processes and templated insertions. Chromothripsis contributes to oncogene amplification as well as to inactivation of genes such as mismatch-repair related genes. These findings show that chromothripsis is a major process driving genome evolution in human cancer.

## INTRODUCTION

Chromothripsis is a mutational phenomenon characterized by massive genomic rearrangements, often generated in a single catastrophic event and localized to isolated chromosomal regions^1–4^. In contrast to the traditional view of tumorigenesis as a gradual Darwinian process of progressive mutation accumulation, chromothripsis provides a mechanism for the rapid accrual of hundreds of rearrangements in few cell divisions.

This phenomenon has been studied in primary tumors of diverse histological origin^5–10^, but similar random joining of chromosomal fragments has also been observed in the germline^11^. There has been considerable progress in elucidating the mechanism by which chromothripsis may arise, including fragmentation and subsequent reassembly of a single chromatid in aberrant nuclear structures called micronuclei^2,12^, as well as fragmentation of dicentric chromosomes in telomere crisis^13,14^. Chromothripsis is not specific to cancer, as it can cause rare congenital human disease and can be transmitted through the germline^11,15^; it has also been described in plants, where it has been linked to micronucleation^16^. However, despite the recent rapid progress on chromothripsis, much remains to be discovered regarding its cause, prevalence, and consequences.

A hallmark of chromothripsis is multiple oscillations between two or three copy number (CN) states^1,6^. Applying this criterion to copy number profiles inferred from SNP arrays, chromothripsis was initially estimated to occur in at least 2-3% of human cancers and in ∼25% of bone cancers^1^. Subsequent studies of large array-based datasets gave similar frequencies: 1.5% (124 of 8,227 tumors across 30 cancer types)^17^ and 5% (918 out of 18,394 tumors across 132 cancer types)^18^, with the highest frequencies detected for soft-tissue tumors (54% for liposarcomas, 24% for fibrosarcomas, and 23% for sarcomas)^18^. These estimates relied on the detection of copy number oscillations that are more densely clustered than expected by chance, *e.g.* at least 10 adjacent CN oscillations in medulloblastomas^8^.

Whole-genome sequencing (WGS) data provide a greatly enhanced view of structural variations (SVs) in the genome with breakpoints identified at single nucleotide resolution. They also provide information on the rearranged DNA sequence, which can be used to determine the type of SVs (*e.g.*, deletion, insertion, inversion) and to infer the likely repair mechanism for joining fragments (*e.g.* based on the degree of microhomology, or the presence or absence of small insertions at the breakpoints). With higher spatial resolution and additional information provided by WGS data, it is possible to postulate a more nuanced set of criteria for chromothripsis and enhance detection specificity^3^. Our earlier analysis of WGS data from cutaneous melanomas already found chromothripsis-like rearrangements in 38% (45 out of 117 patients)^10^; other studies based on WGS found 60-65%^5^ for pancreatic cancer, and 32% for esophageal adenocarcinomas^7^. Whether these examples are outliers reflecting the unique biology of these tumors, or whether they reflect a more general underestimation of the frequency of chromothripsis remained unclear.

Motivated by the importance of chromothripsis during tumor evolution and the need for more systematic and comprehensive analysis, we sought to determine the frequency and spectrum of chromothripsis events in the WGS data for 2,658 cancer patients spanning 39 cancer types, available through the International Cancer Genome Consortium (ICGC). In addition to deriving more accurate per-tumor type prevalence of chromothripsis, we determine the size and genomic distribution of such events, examine their role in amplification of oncogenes or loss of tumor suppressors, describe their relationship to genome ploidy, and investigate whether their presence is correlated with patient survival. Our chromothripsis calls can be browsed at the accompanying website: http://compbio.med.harvard.edu/chromothripsis/.

## RESULTS

### Prevalence of chromothripsis across cancer types

We first sought to formulate a set of criteria for identifying chromothripsis events with varying complexities (Fig. 1a). The generally acknowledged model of chromothripsis posits that some of the DNA fragments generated by the shattering of the DNA are lost; thus, copy number oscillations between two or three states^1,6^ are an obvious first criterion (Fig. 1a). Such deletions also lead to interspersed loss of heterozygosity (LOH), or altered haplotype ratios if there is only a single copy of the parental homolog of the fragmented chromatid. Although chromosome shattering and reassembly has been demonstrated to experimentally generate chromothripsis^2^, template-switching DNA replication errors can generate a similar pattern^19^. Indeed, shattering and replication error models are not exclusive and could indeed co-occur^2^. Therefore, for the discussion below we will refer generally to “chromothripsis” as encompassing both classes of models.

**Figure 1.**
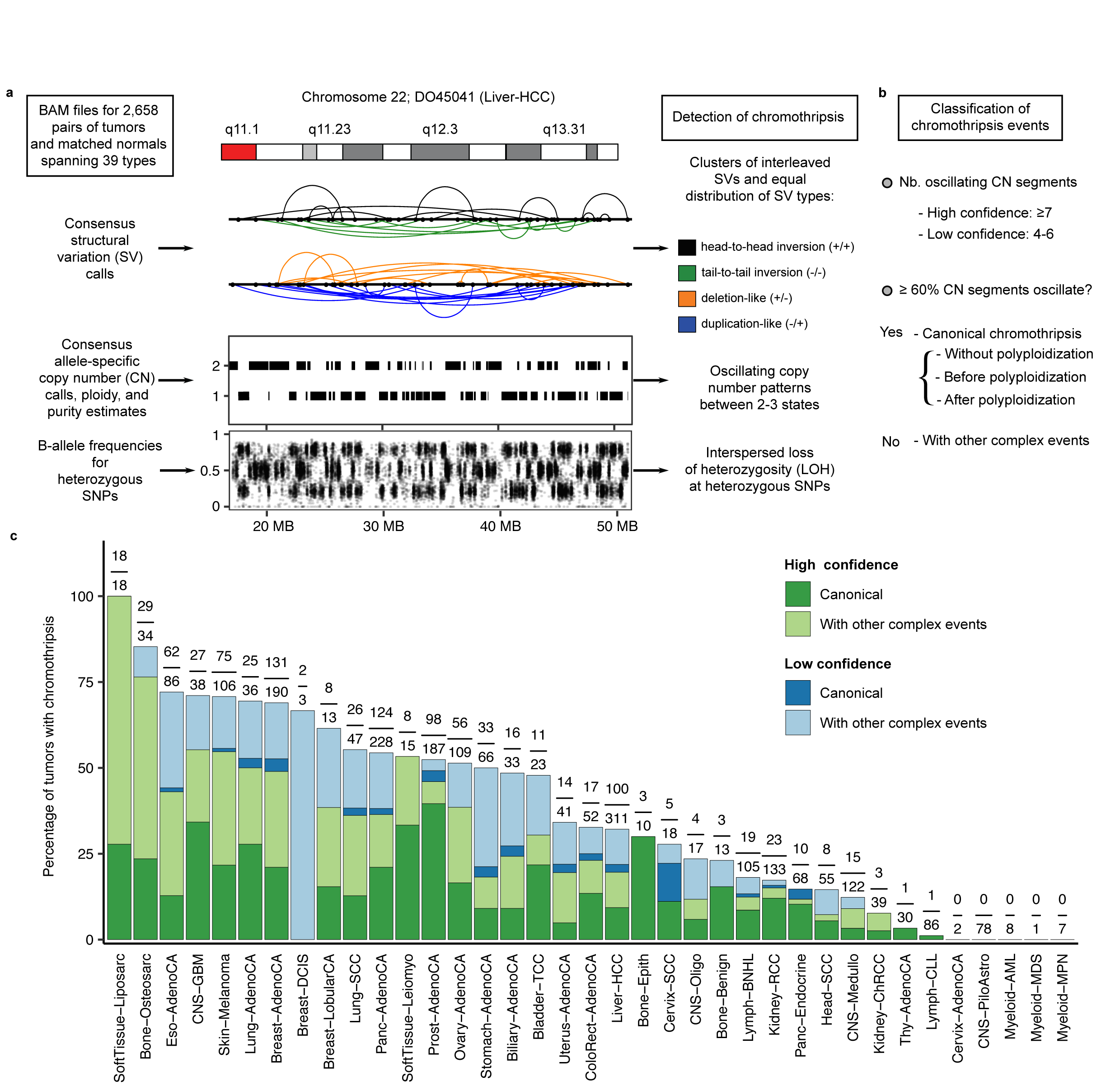
Overview of the chromothripsis calling method and the frequency of events across tumor across 37 cancer types. (**a**) Example of a region displaying the characteristic features of chromothripsis: cluster of interleaved SVs with equal proportion of SV types (*i.e*., fragment joins), CN profile oscillating between 2 states, and interspersed loss of heterozygosity (LOH). Details of the criteria are described in Online Methods. Both the color scheme and the abbreviations shown in this figure are used throughout the manuscript. (**b**) Classification of chromothripsis events. In a canonical event, >60% of the segments oscillate between two CN states; a tumor is classified as canonical if it showed at least one canonical chromothripsis event. (**c**) Percentage of patients harboring chromothripsis events across the entire cohort. The fractions on top of the bars are the number of tumors showing high-confidence chromothripsis over the total number of tumors of that type. The cancer type abbreviations used across the manuscript are as follows: biliary adenocarcinoma (Biliary-AdenoCA), bladder transitional cell carcinoma (Bladder-TCC); bone cartilaginous neoplasm, osteoblastoma, bone osteofibrous dysplasia (Bone-Benign); bone neoplasm, epithelioid (Bone-Epith); sarcoma, bone (Bone-Osteosarc); breast adenocarcinoma (Breast-AdenoCA); breast ductal carcinoma in situ (Breast-DCIS); breast lobular carcinoma (Breast-LobularCA); cervix adenocarcinoma (Cervix-AdenoCA); cervix squamous cell carcinoma (Cervix-SCC); central nervous system (CNS) diffuse glioma (CNS-GBM); CNS diffuse glioma (CNS-Oligo); CNS medulloblastoma (CNS-Medullo); CNS non-diffuse glioma (CNS-PiloAstro); col on/rectum adenocarcinoma (ColoRect-AdenoCA); esophagus adenocarcinoma (Eso-AdenoCA); head/neck squamous cell carcinoma (Head-SCC); kidney chromophobe renal cell carcinoma (Kidney-ChRCC); kidney renal cell carcinoma (Kidney-RCC); liver hepatocellular carcinoma (Liver-HCC); lung adenocarcinoma (Lung-AdenoCA); lung squamous cell carcinoma (Lung-SCC); lymphoid chronic lymphocytic leukemia (Lymph-CLL); lymphoid mature B-cell lymphoma (Lymph-BNHL); lymphoid not otherwise specified (Lymph-NOS); myeloid acute myeloid leukemia (Myeloid-AML); myeloid myelodysplastic syndrome (Myeloid-MDS); myeloid myeloproliferative neoplasm (Myeloid-MPN); ovary adenocarcinoma (Ovary-AdenoCA); pancreatic adenocarcinoma (Panc-AdenoCA); pancreatic neuroendocrine tumor (Panc-Endocrine); prostate adenocarcinoma (Prost-AdenoCA); skin melanoma (Skin-Melanoma) Leiomyosarcoma, soft tissue (SoftTissue-Leiomyo); liposarcoma, soft tissue (SoftTissue-Liposarc) stomach adenocarcinoma (Stomach-AdenoCA) thyroid low-grade adenocarcinoma(Thy-AdenoCA); and uterus adenocarcinoma (Uterus-AdenoCA).

To detect chromothripsis from WGS data, we developed ShatterSeek (a detailed description of the algorithm and its performance is provided in Online Methods and Supplementary Note). A key feature of our method is to identify clusters of breakpoints belonging to SVs that are interleaved, *i.e.*, the regions bridged by their breakpoints overlap instead of being nested (Fig. 1), as is expected from random joining of genomic fragments. This encompasses the many cases that do not display simple oscillations, *e.g.*, partially oscillating CN profiles with interspersed amplifications, and oscillations spanning multiple CN levels due to aneuploidy^5,20^. Rearrangements in chromothripsis should also follow a roughly even distribution for the different types of fragment joins (*i.e*., duplication-like, deletion-like, head-to-head and tail-to-tail inversions, depicted in blue, orange, black, and green, respectively, in Fig. 1a and throughout the manuscript) and have breakpoints randomly distributed across the affected region^1–3^. Finally, by criteria to be described below, we also use interchromosomal SVs to identify chromothripsis events involving multiple chromosomes.

After removing low-quality samples using stringent quality control criteria, we applied our chromothripsis detection method to 2,543 tumor-normal pairs spanning 37 cancer types (Supplementary Table 1 and Online Methods). 2,428 cases harbored SVs and were considered for further analysis. To tune the parameters in our method, we used statistical thresholds and visual inspection. For the minimum number of oscillating CN segments, we used two thresholds: ‘high-confidence’ calls display oscillations between two states in at least 7 adjacent segments, whereas ‘low-confidence’ calls involve between 4 and 6 segments (Fig. 1b and Supplementary Note). The analyses described in the following sections were performed using the high-confidence call set unless otherwise stated.

We first focused on the 1,427 nearly-diploid genomes (ploidy ≤ 2.1; Supplementary Table 1), in which detection of chromothripsis is more straightforward. We defined as ‘canonical’ those events in which >60% of the CN segments in the affected region oscillate between two CN states (canonical events in polyploid tumors are described later). The frequency of canonical chromothripsis events is over 40% for multiple cancer types, such as glioblastomas (CNS-GBM; names in parentheses are short-hand names agreed upon by the ICGC, 50%) and lung adenocarcinomas (Lung-AdenoCA, 40%; Supplementary Note). These numbers provide a lower bounds for the frequency of chromothripsis events, but are nevertheless much higher than previous estimates^17,18^.

When we extend our analysis to the complete tumor cohort (2,543 tumors spanning 37 cancer types that passed our quality-control criteria), we identify high-confidence chromothripsis events in 27% of all the samples (690 out of 2,543), affecting 2.4% of all chromosomes examined (Fig. 1c; Supplementary Data File 1). When low-confidence calls are included, the percentage increases to 38.3% of the samples (975 patients), in 4.1% of chromosomes (Supplementary Data File 2).

The frequency of chromothripsis varies markedly across cancer types. At the high end, we find that 100% of liposarcomas (SoftTissue-Liposarc) and 77% of osteosarcomas (Bone-Osteosarc) exhibit high-confidence chromothripsis (Fig. 1c and Supplementary Fig. 1). Although higher susceptibility of these cancer types to chromothripsis has been described^1,20^, our estimated frequencies are substantially higher. Ovarian adenocarcinomas (Ovary-AdenoCA), breast adenocarcinomas (Breast-AdenoCA), melanomas (Skin-Melanoma), CNS-GBM, esophageal adenocarcinomas (Eso-AdenoCA), and Lung-AdenoCA showed evidence of chromothripsis in >50% of the cases. If we consider low-confidence calls, the frequencies are generally 20-25% higher for these tumors (Fig. 1c). In contrast, the frequencies were lowest in thyroid adenocarcinomas (Thy-AdenoCA; 0%, *n*=30), lymphoid chronic lymphocytic leukemia (Lymph-CLL; 0%, *n*=86), and pilocytic astrocytomas (CNS-PiloAstro; 0%, *n*=78); in the other tumor types with low incidence, the sample sizes are too small to give meaningful estimates.

Overall, these results indicate much greater prevalence of chromothripsis in a majority of human cancers than previously estimated ^10,17,18^. The immense variations we observe across tumor types cannot be explained by random fluctuations and must be linked to tumor biology.

### Understanding the difference between our frequency estimates and previous ones

In accordance with some recent analyses that found dramatically higher frequency of chromothripsis in specific tumor types^5,7^, our overall estimates are considerably higher than those in prior pan-cancer studies. This is in large part due to the fact that previous pan-cancer studies were based on array-based technologies. Additional factors include improvement in SV detection and more refined criteria for defining chromothripsis. In Supplementary Note, we have compiled the criteria used in 26 major chromothripsis-related studies published to date; many did not involve precise description of their approach and code were not publicly available.

To better understand the discrepancy between WGS-based studies, we carried out a detailed comparison on the same datasets that have been analyzed previously. For 109 prostate adenocarcinomas in Fraser *et al*^21^ that were also part of the ICGC dataset, the original authors used ShatterProof^22^—the only publicly available algorithm that uses CNV/SV calls as input—and found chromothripsis in 21% of the tumors (23/109). When we re-applied the same algorithm (with same parameters) but using our CNV/SV calls, the fraction of chromothripsis cases more than doubled to 45% (49/109). This indicates that a major reason for the lower frequencies in the past may be the lower sensitivity in some previous SV detection approaches. SV detection remains a challenging problem especially for low-purity tumors, and algorithms differ substantially in their sensitivity and specificity. Note that the SV calls we used were compiled by the ICGC SV group, with each variant requiring consensus from at least two of four algorithms^23^.

Applying our own chromothripsis algorithm (ShatterSeek), we identified 11 additional cases for a total of 55% (60/109). Compared to the 23 cases reported by Fraser *et al*.^21^, we missed 4. The missed events are focal events comprising less than 6 SVs, which is the lowest number allowed in our criteria to avoid a high false positive rate; the detected regions appeared to be hypermutated regions characterized by tandem duplications or deletions. Visual inspection of the cases we detect but are missed by Fraser *et al*. reveals that the differences in the rates are indeed mostly due to the lower sensitivity of their SV calls (see Supplementary Note for an in-depth comparison). ShatterSeek has increased sensitivity by incorporating cases that display more complex patterns of oscillations and interchromosomal SVs while keeping the specificity high by imposing additional criteria on breakpoint homology to remove tandem duplications and those arising from breakage-fusion-bridge (BFB) cycles. Lastly, we also compared our method against ChromAL^5^ for 76 pancreatic tumors. Both ChromAL and ShatterSeek detect chromothripsis in the same 41 tumors (54%).

Thus, our hypothesis that we have not over-estimated the chromothripsis frequencies is supported by the following: *(i)* some tumor types such as thyroid, CLL, and pilocytic astrocytomas give no events; *(ii)* diploid tumors, which give simpler configurations that are easier to reconstruct or verify visually, give high frequencies; *(iii)* the cases were divided into high- vs low-confidence cases and the high-confidence ones were used for final estimates; *(iv)* more sensitive CNV/SV calls result in higher frequencies for the same datasets; and *(v)* our estimates are in agreement with very recent analysis in specific tumor types. These results reinforce the high prevalence of chromothripsis in human cancers.

### Frequent involvement of interchromosomal SVs

An important feature of our approach is the incorporation of interchromosomal SVs to detect chromothripsis events that involve multiple chromosomes. Chromothripsis affects only a single chromosome in 40.2% of the tumors with chromothripsis (Fig. 2a-c and Supplementary Figs. 1-2). A large number of chromosomes are frequently affected in some tumor types, *e.g*., more than ≥5 chromosomes in 61.1% SoftTissue-Liposarc (Supplementary Figs. 1-2, 3a-d). In one of the most extreme cases, we found a single chromothripsis event affecting six chromosomes (Fig. 2b), where, of the 110 SVs on chromosome 5, only 7 were intrachromosomal. In another example (Supplementary Fig. 3d), a ∼5MB region on chromosome 12 did not display CN oscillations, but it could be linked by interchromosomal SVs to another region that does show a clear chromothripsis pattern, suggesting that the amplification of *CCND2* on chromosome 12 may have originated from chromothripsis. Chromothripsis involving multiple chromosomes can result either from simultaneous fragmentation of multiple chromosomes (*e.g.* multiple chromosomes in a micronucleus or in a chromosome bridge) or from fragmentation of a chromosome that had previously undergone a non-reciprocal translocation.

**Figure 2.**
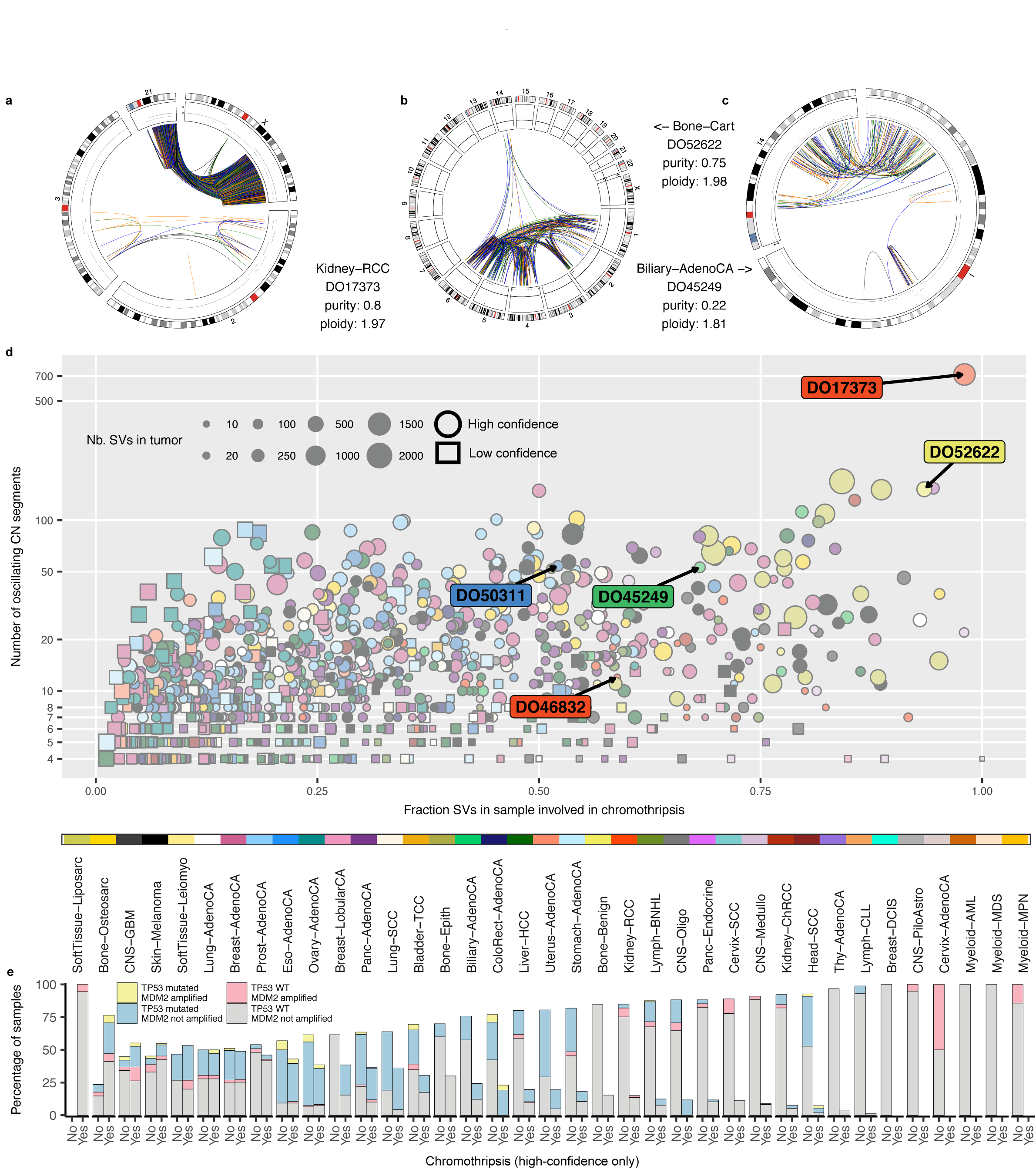
Heterogeneity of chromothripsis events. Examples of massive chromothripsis events on the background of quiescent genomes in patients: DO17373(**a**), DO52622 (**b**), and DO45249 (**c**). (**d**) The fraction of SVs involved in chromothripsis in each sample against the maximum number of contiguous oscillating CN segments for the high (circles) and low-confidence (squares) chromothripsis calls. (**e**) Distribution of patients showing high-confidence chromothripsis, deleterious *TP53* mutations, and *MDM2* amplification (CN≥4).

Distinguishing these scenarios requires a case-by-case analysis that is beyond the scope of this study. However, it is likely that both mechanisms contribute to the generation of chromothripsis involving multiple chromosomes.

### Size and complexity of chromothripsis events are highly variable

Chromothripsis events span a wide range of genomic scale, with the number of breakpoints involved varying by three orders of magnitude within some tumor types (Supplementary Fig. 1c). We find tumors in which relatively focal chromothripsis events, usually a few Mb in size, take place against the backdrop of an otherwise quiet genome (lower-right quadrant in Fig. 2d). Although focal, these events can lead to the simultaneous amplification of multiple oncogenes located in different chromosomes (Supplementary Figs. 3c-e, 4a-c). Other focal events co-localize with other complex events in highly rearranged genomes (lower-left quadrant in Fig. 2d).

Those events spanning large genomic regions comprise tens to hundreds of SVs, affecting anywhere from one chromosome arm to more than 10 chromosomes (Fig. 2a-c). We observe 47 tumors harboring >200 rearrangements, of which at least 50% belong to chromothripsis regions (upper-right quadrant in Fig. 2d). Overall, our analysis reveals greater heterogeneity of chromothripsis patterns than previously appreciated, both in terms of the number of SVs and chromosomes involved.

### Relationship between chromothripsis and aneuploidy

Newly established polyploid cells have high rates of mitotic errors that generate lagging chromosomes^24,25^, which in turn are linked to chromothripsis in medulloblastomas and *in vitro*^2,12,14^. However, a direct causal link between polyploidy and chromothripsis has not been established, and the frequency with which polyploidy and chromothripsis are associated in cancer genomes has not been comprehensively assessed. To examine the sequence of events clearly, we focused on the canonical cases in polyploid tumors, where we can infer whether chromothripsis occurred before or after polyploidization^26^.

For example, if the CN oscillation occurs between 2 and 4 copies in a tetraploid tumor, we infer that polyploidization occurred after chromothripsis; on the other hand, if the oscillation occurs between 3 and 4 copies, we infer that polyploidization occurred first^26^ (Supplementary Figs. 1-2, 4d, 5 and Supplementary Note). Among the canonical cases (∼57% of all chromothripsis events), 68% occur in nearly diploid genomes (n=1,648) and 32% in polyploid tumors (n=748; ploidy >=2.5). Of the 163 cases in which we can distinguish the sequence of events, 74% show chromothripsis after polyploidization while the remaining 26% show chromothripsis before polyploidization. This suggests that a larger fraction of the canonical chromothripsis events in polyploid tumors are late events.

After correcting for tumor type using the logistic regression, we estimate that the odds of chromothripsis occurring in a polyploidy tumor is 1.5 times larger than that in a diploid tumor on average (1.20-1.85 95% CI; P < 10^-3^, cases with ploidy ≥ 2.5). Although polyploidy is associated with higher incidence of chromothripsis, this may be primarily due to the presence of increased genomic material in polyploids. Polyploidy, on the other hand, could reduce the sensitivity of CNV and SV detection (due to lower sequence coverage per copy), and might make it easier for the cell to lose the highly-rearranged copy when intact copies are present^27^.

### Frequent co-localization of chromothripsis with other complex events

About half of the chromothripsis events co-localize with other genomic alterations (Fig. 1 and Supplementary Fig. 1b). Such cases are abundant in SoftTissue-Liposarc (74% of the events), Breast-AdenoCA (65%), Bone-Osteosarc (60%), Skin-Melanoma (59%), Eso-AdenoCA (58%), and Ovary-AdenoCA (55%; Supplementary Figs. 1-2). There is evidence across multiple tumor types that chromothripsis might occur prior or after additional layers of rearrangements^6–8,13,14,28^. For instance, BFB cycles have been postulated to be mechanistically linked to chromothripsis, and telomere attrition, which results in the formation of BFB cycles, has been identified as a predisposing factor for chromothripsis^6,13,29^.

Co-localization of APOBEC-mediated kataegis and rearrangements has been reported for multiple cancer types^30,31^, and has been linked to double-strand break resection during break-induced replication^32^. To study the relationship between kataegis and chromothripsis, we examined the presence of clusters of APOBEC-induced mutations within the chromothripsis regions (Online Methods). Excluding melanoma samples (due to the overlap between the APOBEC and ultraviolet light signatures^33^), we find that 30% of the 690 tumors with chromothripsis show at least 5 clustered APOBEC-induced mutations, and 10% display kataegis comprised of at least 20 mutations. Previous analysis of liposarcomas postulated that multiple BFB cycles on a derivative chromosome generate by chromothripsis underlie the formation of neochromosomes^28^. Consistent with this model, we observe variant allele fractions of 0.01-0.1 for APOBEC-induced SNVs mapped to chromothripsis regions having high-level CN amplifications in Bone-Liposarc tumors, suggesting that they occurred at late stages of tumor development, likely after chromothripsis (Fig. 3e). Overall, although kataegis can co-occur with chromothripsis, this co-occurrence is not common. This is consistent with recent data that chromothriptic derivative chromosomes are mostly assembled by end joining mechanisms which do not involve extensive DNA end resection^34^.

### *TP53* mutation status and chromothripsis

Inactivating *TP53* mutations have been previously associated with chromothripsis in medulloblastomas^8^ and in pediatric cancers^35,36^. *TP53* deficient cells serve as models to generate chromothripsis *in vitro*^2,14^. Nevertheless, the relationship between deleterious *TP53* mutations and chromothripsis has not been examined comprehensively. In our data, ∼38% of the patients with inactivating *TP53* mutations show chromothripsis, whereas only 24% of those with wildtype *TP53* exhibit chromothripsis (Fig. 2e). After correcting for cancer type, this translates to an odds ratio of 1.54 (1.21-195, 95% CI, *P*<10^-3^) for chromothripsis in patients with *TP53* mutations compared to *TP53* WT cases, reinforcing the notion that *TP53* mutations are associated with a higher incidence of chromothripsis. However, we note that 398 (58%) of the patients in our cohort exhibiting chromothripsis do not show *TP53* mutations nor *MDM2* amplifications (the major cellular regulator of *TP53* by ubiquitination^37^), including those with massive cases in diploid genomes, *e.g.*, DO25622 (Fig. 2b). This suggests that, although *TP53* malfunction and polyploidy are predisposing factors to chromothripsis, it still occurs frequently in diploid tumors with proficient *TP53*.

### Signatures of repair mechanisms in chromothripsis regions

By examining the sequence homology at the breakpoints, it is possible to infer the predominant mechanisms responsible for the chromothripsis event^38,39^. Although this classification is not precise, it is helpful in providing an overview of mutational signatures. Previously, NHEJ has been implicated in the reassembly of the DNA fragments generated by chromothripsis^2,34^, whereas alternative end-joining (alt-EJ) has been proposed to occur in constitutional chromothripsis and in glioblastomas^15,40^. In addition, short templated insertions suggestive of microhomology-mediated break-induced replication (MMBIR) or alt-NHEJ associated with polymerase theta have been detected in chromothripsis originated from DNA fragmentation in micronuclei^2,41–43^.

We analyzed the breakpoints involved in canonical chromothripsis events showing evidence of interspersed LOH, as most SVs in such cases are chromothripsis-related (Fig. 1b). In 55% of these events, we only detected repair signatures concordant with NHEJ or alt-EJ (Supplementary Fig. 6). In 32%, we identified stretches of microhomology at two or more breakpoint junctions (most of them of 0-6 bp) and short insertions of 10-500 bp that map to distant locations within the affected region by chromothripsis (Supplementary Fig. 6). For instance, in the massive chromothripsis case depicted in Fig. 2a (where we detect 1,962 SVs, hundreds of uninterrupted CN oscillations between CN states 1 and 2, and interspersed LOH), we detect small nonrandom insertions of 10-379 bp at 60 breakpoints. Thus, NHEJ plays a principal role in DNA repair, with partial contribution of MMBIR or alt-NHEJ in generating canonical chromothripsis.

By contrast, ∼5% of the canonical events detected in diploid genomes show no evidence of LOH in part or the entire region affected (*e.g*., oscillations between 2 and 3 CN, long stretches of microhomology, and frequent evidence of template switching (Figs. 3 and 4)^23^. For instance, in the case depicted in Fig. 3b affecting chromosome 4 of an Ovary-AdenoCA tumor, both the size of the segments at CN 3 (*i.e*., in the 3-285Kb range; mean of 45Kb) and the orientation of the breakpoints at their edges (*i.e*, − and +), suggest that these are templated insertions^23^. In addition, multiple breakpoint junctions show features concordant with MMBIR. For this particular case, we could manually reconstruct part of the amplicon by following the polymerase trajectory across 43 template switching events (Fig. 3c-f). This type of event might be more appropriately called chromoanasynthesis^19^, but systematically distinguishing chromoanasynthesis from chromothripsis in cancer genomes is currently challenging due to their partially overlapping features (template switching events can generate LOH if the polymerase skips over segments of the template, and LOH might not be present in chromothripsis events occurring in aneuploid genomes; Supplementary Note).

**Figure 3.**
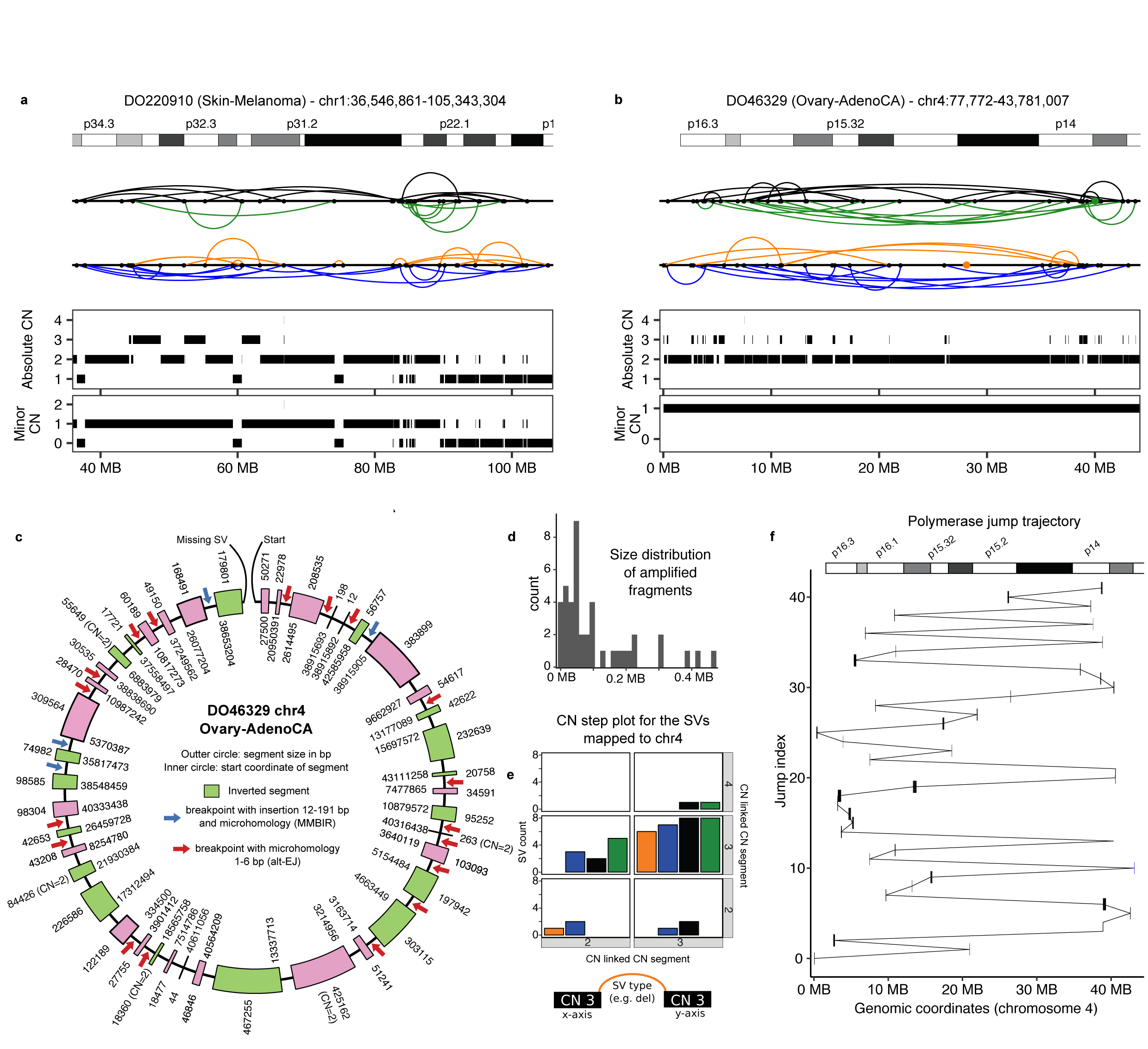
Example of canonical chromothripsis events displaying templated insertions and evidence of MMBIR. (**a**) Evidence of chromothripsis in chromosome 1 in a Skin-Melanoma tumor displaying CN oscillations spanning 3 CN levels and LOH. (**b**) Example of a chromothripsis event in chromosome 4 involving low-level copy gains and absence of LOH in an Ovary-AdenoCA tumor. Segments at CN 3 correspond to templated insertions, as evidenced by their size, and breakpoint orientations at their edges. Breakpoints corresponding to interchromosomal SVs are depicted as colored dots in the SV profile, whereas intrachromosomal SVs are represented with black dots and colored arcs following the representation shown in Fig. 1. (**c**) Reconstruction of the amplicon generated by the chromoanasynthesis event detected in chromosome 4 in tumor DO46329 (see **b**). Inverted segments are depicted in green. Red arrows highlight breakpoints with short microhomology tracts, whereas blue arrows indicate the presence of small insertions at the breakpoints. The CN for all segments is 3 unless otherwise indicated. (**d**) Size distribution for the templated insertions forming the amplicon depicted in **c**. (**e**) CN step plot for chromosome 4 indicating that most of the SVs mapped to chromosome 4 link genomic regions at CN 3. The x and y axes correspond to the CN level of the segments linked by a given SV. The colour of the bars correspond to the four types of SVs (i.e. dup-like, del-like, and inversions) indicated in Fig. 1a and considered throughout the manuscript. (**f**) Trajectory of the polymerase across chromosome 4 estimated from the template switching events depicted in **c**.

**Figure 4.**
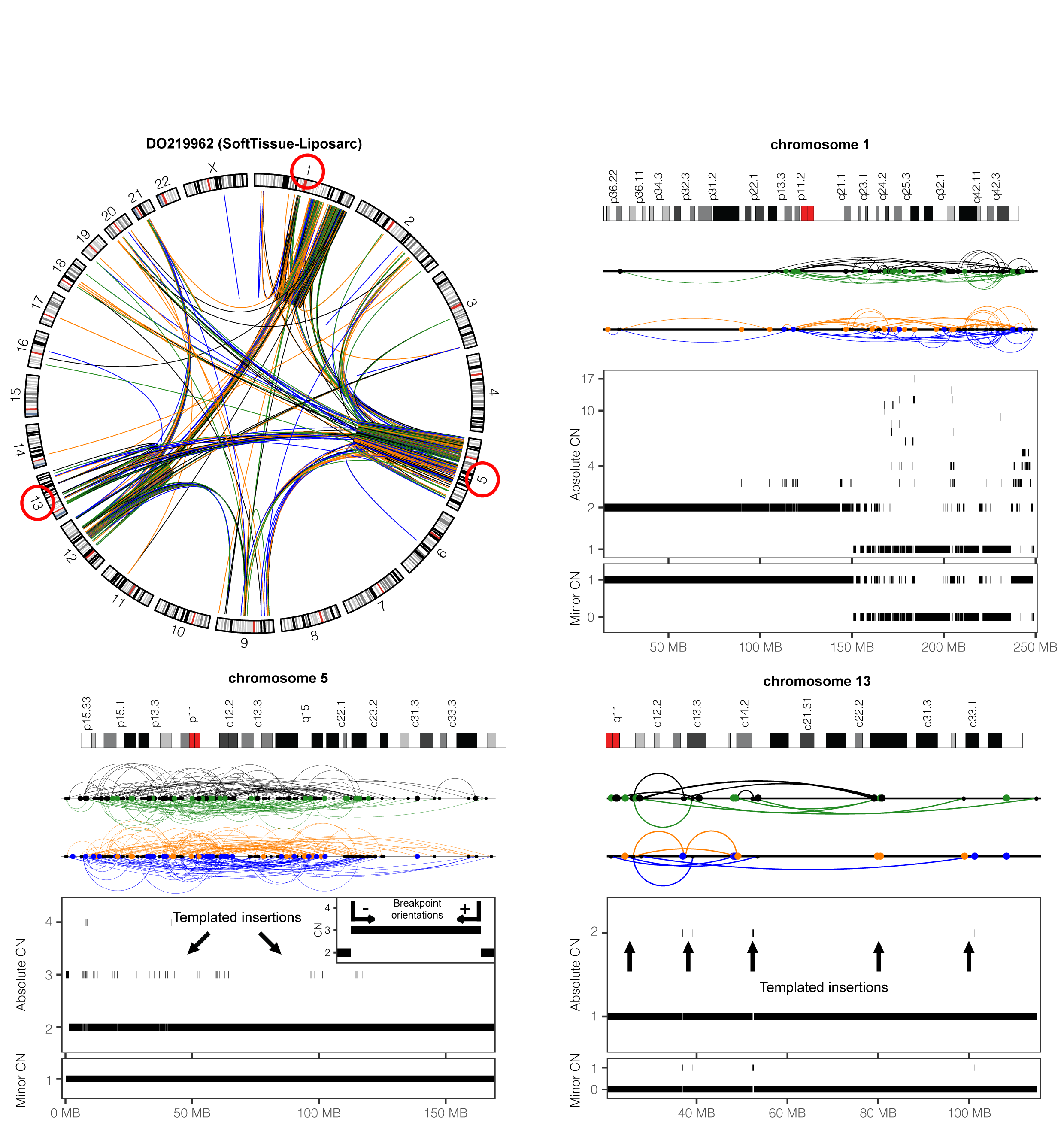
Example of a multichromosomal chromothripsis event in a SoftTissue-Liposarc tumor co-localized with other complex events involving templated insertions. (**a**) Scaled circos plot of the entire genome for this tumor except for chromosome Y. (**b-d**) SV and CN profiles for chromosomes 1 (**b**), 5 (**c**) and 13 (**d**). Tens of CN oscillations and LOH in chromosome 1 are co-localized with additional rearrangements. The size, minor CN, and the orientation of the breakpoint junctions associated to the segments at CN 3 indicate that these are templated insertions. The inset in (**c**) represents the orientation of the breakpoint junctions at the edges of low-level CN gains originated from template switching (*i.e.*, and + according to the notation we use in the manuscript).

We also find features associated to replication-associated mechanisms in more complex rearrangements involving multiple chromosomes (Fig. 4). In a number of these events, LOH is observed in some chromosomes (*e.g*., Fig. 4b), but it is absent in others, where the oscillations occur at higher CN states without LOH (Fig. 4c,d). For instance, in the case reported in Fig. 4, there is evidence of templated insertions in chromosomes 5 and 13, which are linked to a chromothripsis event showing LOH in chromosome 1. Notably, the minor CN (*i.e*., the copy number of the allele with the lower number of copies) for the templated insertions in chromosome 13 is 1, whereas for the rest of the chromosome is 0. This suggests that one parental chromosome served as a template and was later lost to the cell.

Overall, these results hint at the involvement of template switching events in the generation or repair of complex rearrangements, consistent with the observations of replicative processes in the formation of clustered rearrangements in congenital disorders and cancer^15,19,38,23,44^. Although further experimental evidence will be necessary, we postulate that the involvement of replication-associated mechanisms in the assembly of derivative chromosomes in chromothripsis might be substantial.

### Oncogene amplification in chromothripsis regions

Evidence of double minute (DM) formation from chromothripsis has been reported for selected cancer types^1,2,8,40^. However, the extent to which chromothripsis contributes to DM formation has not been fully examined on a pan-cancer scale. Although reconstruction of DM structure with appropriate discordant reads would present clear evidence for its extrachromosomal nature, this proves to be too difficult in most cases. Therefore, we rely on copy number to make our inferences. We find that 15 patients (2% of tumors with chromothripsis) show CN oscillations between one low (CN≤4) and one very high (CN≥10) states, consistent with the presence of DM^8,40^. We detect known cancer drivers in these putative DMs, including *MDM2* (4 samples, Supplementary Figs. 3e and 4a, and Supplementary Table 2), and *CDK4* (4). These amplifications lead to increased mRNA levels: *e.g.*, 5 log_2_-fold increase for *MDM2, NUP107*, and *CDK4* in a GBM sample (DO14049) compared to other GBM tumors. In chromothripsis regions subject to additional rearrangements, it is not always possible to discern using bulk sequencing data alone whether highly amplified segments are part of DMs or correspond to intrachromosomal gene amplification^45^. Furthermore, once a DM has formed, the derivative chromosome showing chromothripsis may be lost if it has no other tumor-promoting mutations. Therefore, the contribution of chromothripsis to the formation of extrachromosomal DNA bodies is likely to be higher than estimated here.

Further analysis of focal amplifications, defined as regions at CN ≥ 4 and smaller than 6Mb^46^, in 1,268 tumors and 162 normal tissue samples with RNA-seq data available reveals that 6,310 focal amplifications encompassing oncogenes (11.1%; 20.5% when including low-confidence calls) localize to chromothripsis regions, and often leading to increased expression (Supplementary Table 2). These include well-known cancer genes, such as *CCND1* (25 tumors), *CDK4* (25), *MDM2* (23), *SETDB1* (23), *ERBB3* (11), *ERBB2* (11), *MYC* (10), and *MYCN* (5). Thus, chromothripsis, perhaps with associated replication-based copy number gains^20,47^, may make a significant contribution to small-scale focal amplifications.

### Chromothripsis-mediated loss of tumor suppressors and DNA repair genes

Tumor suppressors have a direct effect on cell growth or genes, such as those involved in DNA repair, or accelerate the rate of acquiring other growth-promoting or survival mutations^48^. Except for few selected cancer types^5,49^, the extent to which chromothripsis contributes to the loss of these genes has not been thoroughly examined yet. We find that chromothripsis underlies 2.1% and 1.9% of the losses of tumor suppressors and DNA repair genes, respectively. These include *MLH1* (9 patients out of 301 harboring *MLH1* deletions), *PTEN* (12/358), *BRCA1* (8/154), *BRCA2* (7/270), *APC* (9/201), *SMAD4* (10/403), and *TP53* (8/614), among others (Supplementary Fig. 7 and Supplementary Table 2). In 28 of these cases (28 genes in 24 tumors), both alleles were inactivated, one due to chromothripsis and the other due to an SNV. These include *SMAD4, APC, TP53*, and *CDKN2A*. In a biliary adenocarcinoma (Fig. 5), for instance, one *MLH1* allele was lost due to chromothripsis and the other allele was likely silenced due to promoter hypermethylation, as evidenced by low expression of *MLH1* and the microsatellite instability phenotype in an otherwise MMR proficient tumor^50^. Overall, these data illustrate the way in which chromothripsis can confer tumorigenic potential through the loss key tumor suppressors and DNA repair genes.

**Figure 5.**
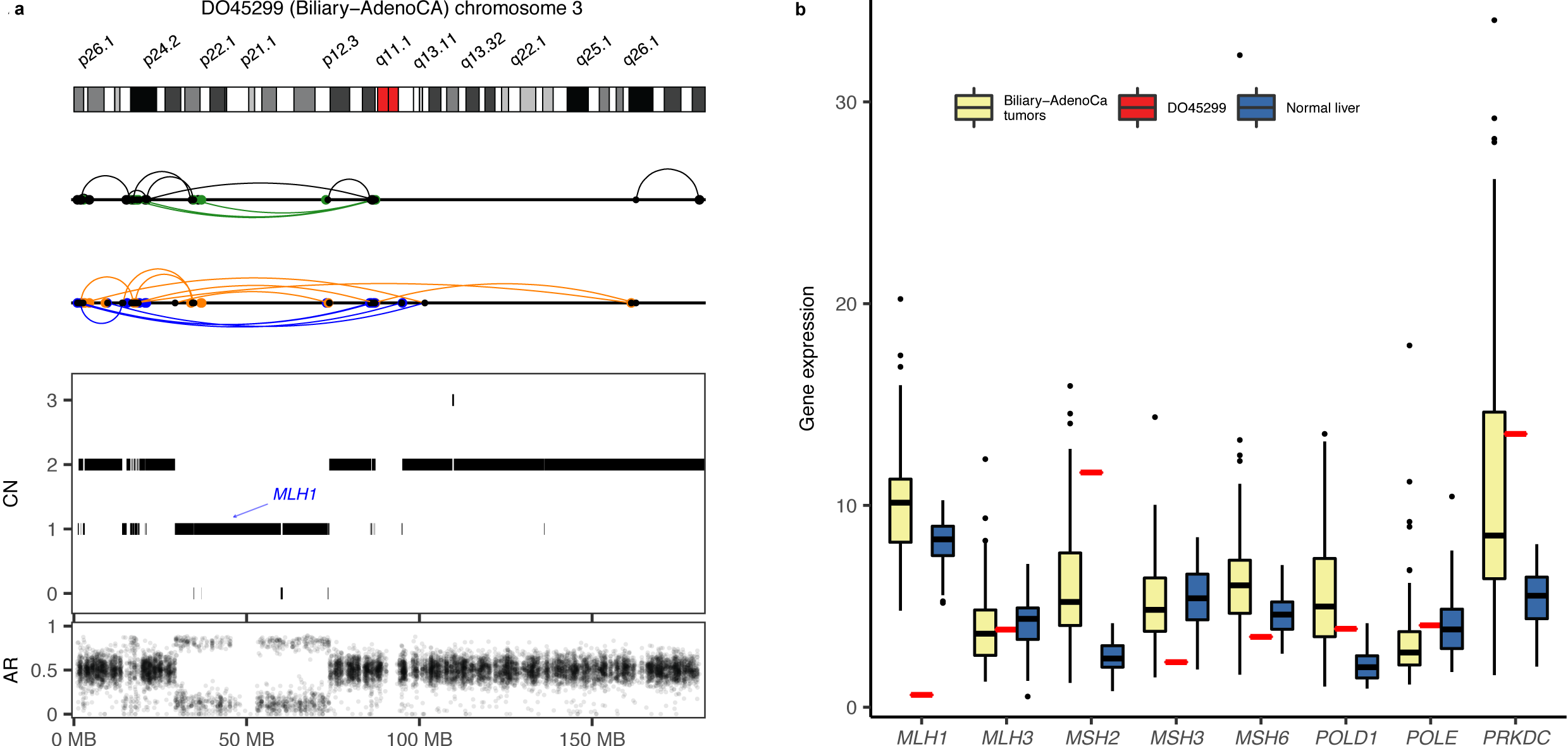
Chromothripsis-mediated depletion of *MLH1*. (**a**) Chromothripsis event and expression levels of DNA mismatch repair (MMR) genes in patient DO45299 (Biliary-AdenoCA), (**b**) their mean expression in a panel of 16 Biliary-AdenoCA tumors, and 16 normal liver samples.

### Chromothripsis is prognostic of poor patient survival

Chromothripsis has been associated with poor prognosis for several cancer types^5,8,9,51^. Here, the increased sensitivity of our approach and the larger cohort permits us to evaluate the impact of chromothripsis on patient survival in greater detail, and to determine the biological contexts in which chromothripsis leads to more aggressive tumors. We did not find significant association between chromothripsis and survival in a multivariate pan-cancer analysis when stratifying the patients into two categories with respect to the presence or absence of chromothripsis (Wald and likelihood ratio tests (LRT)<10^-15^; hazard ratio (HR): 1.07, *P*=0.24). However, we find a significant association in a stratified analysis when the samples are ordered by the percentage of SVs involved in chromothripsis in each tumor type and classified into three equal-sized groups ‘absent’, ‘moderate’, and ‘predominant’ (Wald and LTR<10^-15^; Supplementary Fig. 8a).

The HR for the samples in the predominant and moderate chromothripsis categories are 1.28 (*P*<0.001) and 1.11 (*P*=0.12), respectively. This trend is also observed when also including the low-confidence calls in the analysis, namely 1.26 (*P*<0.001) and 1.12 (*P*=0.10), respectively.

Several factors could influence whether chromothripsis is more prevalent in diploid or polyploid tumors. In a cancer primarily driven by tumor suppressor loss, the growth-promoting effects should be more penetrant in a diploid tumor relative to a polyploid tumor if chromothripsis happened after the genome doubling. To address the relationship between chromothripsis and polyploidy, we divided the patients into 6 categories depending on the penetrance of chromothripsis (*i.e.* absent, moderate and predominant) and ploidy, *i.e.* diploids *vs* polyploids (ploidy ≥ 2.5). We find that the association between chromothripsis and poor prognosis is stronger when it occurs in diploid tumors harboring either moderate (HR_diploids_: 1.18, *P*=0.04; HR_polyploids_: 0.96, *P*=0.68) or predominant chromothripsis (HR_diploids_: 1.30, *P*=0.002; HR_polyploids_: 1.21, *P*=0.06; Supplementary Fig. 8b).

## DISCUSSION

Our analysis of ca. 2,600 cancer genomes has revealed that chromothripsis plays a major role in shaping the architecture of cancer genomes across diverse human cancers, with prevalence and heterogeneity much higher than previously appreciated and marked variability across cancer types. Our approach enabled us to define more nuanced criteria to detect chromothripsis events, including those that involve multiple chromosomes and those that were hard to detect previously due to the presence of other co-localized rearrangements.

We note that the estimated frequencies of chromothripsis depend on the cut-off values used for statistical significance. We have tested various parameters and chose conservative thresholds, such as at least 7 CN segments oscillating between two copy number states for the high-confidence calls; however, we cannot exclude the possibility that some of these chromothripsis-like patterns might have arisen due to other sources of genomic instability. Conversely, it is also possible that we missed true chromothripsis events that have fewer than the required number of rearrangements; it is worth noting that such small-scale events are seen in experimentally generated chromothripsis^2^.

Cases in which chromothripsis is followed by other complex rearrangements that mask the canonical CN pattern are especially difficult to detect, requiring additional criteria and in-depth manual inspection. Despite these limitations, we believe that our statistical approach based on observed frequencies of various alterations compared to the background is more sensitive than a reassembly-based approach. The latter method attempts to reconstruct the steps that led to the observed SV pattern, but most complex events are too complicated, especially when many breakpoints are entirely missed and some are incorrectly identified due to inherent limitation of short-read data, imperfect SV algorithms, and insufficient sequencing coverage.

A substantial fraction of the chromothripsis events we detect show templated insertions and evidence of MMBIR. Although chromothripsis and chromoanasynthesis are considered to be two different processes, they often lead to similar SV and CN profiles, especially when occurring in aneuploid genomes, as oscillating CN profiles with interspersed LOH might be generated by replicative processes alone if the DNA polymerase skips over segments of the template. Moreover, there is experimental evidence that MMBIR and NHEJ can co-exist in chromothripsis induced in micronuclei^2^. Therefore, further experiments will be required to assess the interplay between DNA repair mechanisms in chromothripsis.

Given the pervasiveness of chromothripsis in human cancers and its association with poorer prognosis, another question that arises is whether chromothripsis *per se* constitutes an actionable molecular event amenable to therapy. This is particularly interesting given the link between aneuploidy, depleted immune infiltration, and reduced response to immunotherapy^52^. As more WGS data are linked to other data types including clinical information, it will become more feasible to understand the impact of chromothripsis on tumorigenesis and its potential as a biomarker for diagnosis or treatment.

## Online Methods

### Whole-genome sequencing data

We integrated in a common processing pipeline whole-genome sequencing data from the TCGA and ICGC consortia for 2,658 tumor and matched normal pairs across 39 cancer types, of which 2,543 pairs spanning 37 cancer types that passed our quality-control criteria were selected for further analysis ^53^. The list of samples is provided in Supplementary Table 1. Further information for all tumor samples and patients is given in54. Sequencing reads were aligned using BWA-MEM 0.7.8-r455, whereas biobambam 0.0.138 was used to extract unpaired reads and mark duplicates^55,56^.

### Mutation calling

We utilized the consensus SNV and indel calls sets released by the Pan-Cancer Analysis of Whole Genomes (PCAWG) project, whereas we used HaplotypeCaller 3.4-46-gbc0262554 to call single nucleotide polymorphisms (SNPs) in both tumor and matched normal samples following the GATK best practices guidelines. We only kept SNPs supported by at least 10 reads. We processed a total of 210,021 non-synonymous somatic mutations, of which 43,548 were predicted as deleterious using the MetaLR score as implemented in Annovar ^57^. To identify APOBEC mutagenesis we followed the procedure previously described ^33^. In brief, we considered as APOBEC-associated mutations those involving a change of (i) G within the sequence motif wGa to a C or A (where w is A or T), and (ii) C in the sequence motif tCw to G or T (where w is A or T).

### Detection of SVs and SCNAs

The SVs were identified by the ICGC SV subgroup, which applied four algorithms and selected those SVs found by at least two algorithms ^23,54^. We used the consensus SV, SCNA, purity and ploidy call sets generated by the PCAWG project. The calling pipelines are described in detail in the accompanying manuscripts [(i) Dentro, Leshchiner, Haase, Wintersinger, *et al.* Pervasive intra-tumour heterogeneity and subclonal selection across cancer types; (ii) PCAWG-6. Signatures of selection for somatic rearrangements across 2,693 cancer genomes].

### RNA-seq data analysis

We processed RNA-seq data for a total of 162 and 1,268 normal and tumor samples, respectively. Sequencing reads were aligned using TopHat2 and STAR ^58,59^. HTseq-count was subsequently used to calculate read counts for the genes encompassed in the PCAWG reference GTF set, namely Gencode v19. Counts were normalized to UQ-FPKM (upper quartile normalized fragments per kilobase per million mapped reads) values using upper quartile normalization. The expression values were averaged across the two alignments. The set of oncogenes was downloaded and curated from COSMIC (dominant genes) and IntOGen databases ^60,61^, whereas the set of tumor suppressors was downloaded from TSGene 2.0, COSMIC (recessive genes) and ^62,63^. DNA repair genes were extracted from ^64^.

### Characterization of chromothripsis events using ShatterSeek

To identify and visualize chromothripsis-like patterns in the cancer genomes using CN and SV data, we extended the set of statistical criteria proposed by ^3^ into a calling pipeline composed of the following steps. The ShatterSeek code, the package documentation, and a detailed tutorial are available at https://github.com/parklab/ShatterSeek. Interactive Circos plots for all tumors in PCAWG are provided at http://compbio.med.harvard.edu/chromothripsis/.

The values for the statistical criteria for all chromosomes across all samples are given in Supplementary Table 1. Visual depictions of the high-confidence and low-confidence calls are given in Supplementary Data Files 1 and 2. Visual depictions for the two sets of SV clusters not identified as chromothripsis by our method, namely: (i) those involving clusters of duplications or deletions leading to copy number oscillations, as well as oscillating CN profiles with few or no SVs mapped; and (ii) large clusters of interleaved SVs not displaying chromothripsis are given in Supplementary Data Files 3 and 4, respectively. In all Supplementary Data Files and in the main text, intrachromosomal SVs are depicted as arcs with the breakpoints represented by black points, whereas the breakpoints corresponding to interchromosomal SVs are depicted as colored points.

Duplication-like SVs, deletion-like SVs, head-to-head and tail-to-tail inversions are depicted in blue, orange, black, and green, respectively. The value for the statistical criteria described above for each event is provided below its representation.

### Survival analysis

We performed multivariate analysis using the Cox proportional hazards model corrected for confounding factors known to influence survival rates, namely: age at the time of diagnosis, sex, tumor stage, radiation therapy, the presence of metastasis, and cancer type. Survival analysis was conducted using the *coxph* function from the R package survival version 2.30. Significance was assessed by the likelihood ratio and Wald tests. Tumor stages for all cancer types were manually curated and grouped into four categories. In the pancancer analysis, we considered only cancer types with clinical data available for at least 20 patients. The clinical data are listed in Supplementary Table 1.

## Data availability

The code for calling chromothripsis events is available at https://github.com/parklab/ShatterSeek.

### Acknowledgements

The results published here are partly based upon data generated by The Cancer Genome Atlas and obtained from the Database of Genotypes and Phenotypes (dbGaP) with accession number phs000178.v8.p7. Information about TCGA can be found at http://cancergenome.nih.gov. We thank the Research Information Technology Group at Harvard Medical School for providing computational resources. This project has received funding from the European Union’s Framework Programme For Research and Innovation Horizon 2020 (2014-2020) under the Marie Curie Sklodowska-Curie Grant Agreement No. 703543 (I.C.C.). This work was supported by grants from the Ludwig Center at Harvard (I.C.C., J.K.L. and P.J.P.). We thank Scott Ouellette for his help in deploying the companion website.

## Author contributions

I.C.C. performed bioinformatic analysis of all data, supervised by P.J.P. R.X. developed the initial version of the chromothripsis detection pipeline. D. J. and Y.L.J. contributed bioinformatics analysis of gene expression data. The manuscript was written by I.C.C., J.K.L. and P.J.P, with substantial input from C.-Z.Z. and D.S.P. D.M. and L.J.K. performed analysis of APOBEC-associated mutations. All authors read and approved the final version of this manuscript.

## Supplementary Figures

**Supplementary Figure 1.** Landscape of chromothripsis across 37 cancer types including high-confidence calls only. (**a**) Percentage of patients harboring chromothripsis events. The total number of samples examined from each cancer type is indicated on top of the bars, whereas the colors of the bars indicate the number of chromosomes affected by chromothripsis. (**b**) Number of chromothripsis regions categorized on the basis of the oscillating copy number states and the temporal profile of the events with respect to polyploidization, and their distribution across the genome. (**c**) Distribution of the number of breakpoints (blue), and span of the chromothripsis events in megabases (pink).

**Supplementary Figure 2.** Landscape of chromothripsis across 37 cancer types including the high- and low-confidence calls. (**a**) Percentage of patients harboring chromothripsis events. The total number of samples examined from each cancer type is indicated on top of the bars, whereas the colors of the bars indicate the number of chromosomes affected by chromothripsis. (**b**) Number of chromothripsis regions categorized on the basis of the oscillating copy number states and the temporal profile of the events with respect to polyploidization, and their distribution across the genome. (**c**) Distribution of the number of breakpoints (blue), and span of the chromothripsis events in megabases (pink).

**Supplementary Figure 3.** (**a**) Number of chromothripsis events involving two given chromosomes across the entire set of high-confidence calls. (**b**) Unscaled circos plot showing a massive chromothripsis event in patient DO52617 (Bone-Osteosarc) involving 16 chromosomes. (**c**) Example of a predominant chromothripsis event in patient DO35845 (CNS-Medullo) affecting a few megabases in chromosomes 3 and 8, with an oscillating copy number pattern between states 4 and 49. This case also illustrates the amplification of *MYC*. Panel (**a**) shows the SVs detected across the entire genome, whereas panel (**b**) corresponds to the regions affected by chromothripsis (scaled circos plot). (**d**) Scaled circos plot representing the chromothripsis event in patient DO13971 (TCGA-06-0221; CNS-GBM) involving chromosomes 2 and 12, leading to the concomitant amplification of *MYCN* and *CCND2*. (**e**) Concomitant amplification of *CDK4* and *MDM2* in chromosome 12 in patient DO219966 (TCGA-DX-A3LT; Bone-Leiomyo).

The bottom panel represents the variant allele fraction (VAF) for somatic SNVs. APOBEC-associated mutations are colored in red.

**Supplementary Figure 4.** (**a**) Example of a focal chromothripsis event in patient DO14049 (TCGA-19-2624; CNS-GBM) involving chromosomes 7 and 12 leading to the concomitant amplification of *CDK4, MDM2*, and *EGFR*. A previous study where this tumor was studied using single-cell sequencing reported that heterogeneity in the *EGFR* locus defined different subclones^45^. These results are concordant with our calls in that the chromothripsis event affecting the *EGFR* locus presents CN oscillations across two states in the regions containing *MDM2* and *CDK4*, whereas the oscillating pattern is distorted at the *EGFR* locus, consistent with the presence of secondary subclonal rearrangements. The lack of a clear oscillation between two CN states would be expected from bulk sequencing data if secondary rearrangements had further altered the *EGFR* locus in different subclones. Although the canonical pattern of chromothripsis is not fully apparent in the *EGFR* locus when using bulk data, we could correctly detect chromothripsis at that locus by using interchromosomal SVs. (**b**) Copy-number step plot6. Each cell represents the distribution of types (see Fig. 1C) for those SVs connecting segments of the CN equal to the values indicated in the *x* and *y*-axis. (**c**). Example of a chromothripsis event in chromosome 12 in patient DO14049 (TCGA-19-2624; CNS-GBM) leading to the concomitant amplification of *CDK4, MDM2*, and *NUP107* likely as a result of DM formation (see also Supplementary Fig. 7).

(**d**) Distribution of the two first CN modes for the chromothripsis events categorized as canonical without polyploidization (black), canonical after polyploidization (green), canonical before polyploidization (red), and chromothripsis events in the context of other complex events (gray). Cases with CN modes higher than 10 were not depicted for clarity.

**Supplementary Figure 5.** Examples of chromothripsis regions. (**a**) canonical chromothripsis without polyploidization (chromosome 22; DO45041, Liver-HCC), (**b**) canonical without polyploidization displaying APOBEC-mediated kataegis (chromosome 4; DO52615, Bone-Osteosarc), (**c**) canonical chromothripsis after polyploidization (chromosome 2; DO52620, Bone-Osteosarc), and (**d**) canonical chromothripsis before polyploidization (chromosome 12; DO36459, ColoRect-AdenoCA). Clustered APOBEC-associated mutations are shown as red dots.

**Supplementary Figure 6.** Analysis of the DNA repair mechanisms for the high-confidence chromothripsis events. (**a,b**) Distribution of the length of the insertions and homology sequences, respectively, detected at the breakpoints for the high-confidence cases with at least 80% of the breakpoints detected pertaining to chromothripsis (predominant chromothripsis). Percentage of breakpoints for each DNA repair mechanism across all cases displaying canonical chromothripsis without polyploidization (**c**), canonical chromothripsis after polyploidization (**d**), canonical chromothripsis before polyploidization (**e**), and chromothripsis co-localized with other complex rearrangements (**f**).

**Supplementary Figure 7.** Chromothripsis-mediated loss of tumor suppressors. (**a**) Evidence of chromothripsis-mediated loss of one *TP53* allele in patient DO11148 (TCGA-14-1034; CNS-GBM). The remaining allele harbors a deleterious mutation (pathogenic missense SNV, p.R337C; COSM11071). (**b**) Chromothripsis-mediated loss of one *SMAD4* allele in patient DO38388 (TCGA-BR-7722; Stomach-AdenoCA). The remaining allele harbors a deleterious mutation (pathogenic missense SNV, chr18: 48,591,925 G>A). (**c**) Biallelic *BRCA2* deletion in patient DO4080 (Breast-AdenoCA) confirmed by the low expression levels of *BRCA2*). This tumor also exhibited the mutational signature 3, which is known to be related to *BRCA1*/*BRCA2* deficiency. (**d**) Monoallelic loss of *APC* in patient DO44883 (Liver-HCC).

**Supplementary Figure 8.** Association between chromothripsis and patients survival. Samples were stratified into three categories based on the fraction of SVs that map to chromothripsis regions: absent (black; no chromothripsis), moderate (“mod.”; red; the fraction of chromothripsis-related SVs is smaller than the median within the same cancer type), and predominant (“pred.”; blue; the fraction of chromothripsis-related SVs is higher than the median within the same cancer type). Kaplan-Meier plots and estimated Hazard Ratios (HR; 95% confidence intervals) for (**a**) all samples stratified on the basis of chromothripsis. Even after accounting for other variables (Cox model), the predominantly chromothripsis samples have significantly worse survival than those without chromothripsis (HR=1.26, *P*=0.001). (**b**) chromothripsis and ploidy. For the diploid samples, the negative impact of chromothripsis is stronger, with both pred and mod groups reaching statistical significance (*cf*. (**a**)).

## Supplementary Tables

**Supplementary Table 1.** The results for the statistical criteria implemented to detect chromothripsis for the 2,428 patients harboring structural variations.

**Supplementary Table 2. (a)** Genes detected in focally amplified regions. (**b**) Mutated genes in focally amplified regions. (**c**) Genes detected in chromothripsis regions at CN 1 or 0. (**d**) Genes with one allele lost due to chromothripsis, and the other harboring deleterious mutations.

## Supplementary Data Files

**Supplementary Data File 1.** High-confidence chromothripsis calls.

**Supplementary Data File 2.** Low-confidence chromothripsis calls.

**Supplementary Data File 3.** Regions displaying CN oscillations not classified as chromothripsis. These include: (i) CN oscillating profiles characterized by clusters of tandem duplications or deletions, (ii) candidate chromothripsis cases satisfying the statistical criteria but considered false positives by visual inspection, and (iii) chromosomes displaying at least 7 CN oscillations with few or no SVs mapped.

**Supplementary Data File 4.** Large clusters of interleaved SVs (>20) not identified as chromothripsis by our method.

